# Reward Evokes Visual Perceptual Learning Following Reinforcement Learning Rules

**DOI:** 10.1101/760017

**Authors:** Zhiyan Wang, Dongho Kim, Giorgia Pedroncelli, Yuka Sasaki, Takeo Watanabe

## Abstract

Visual perceptual learning (VPL) is defined as a long-term performance enhancement as a result of visual experiences. A number of studies have demonstrated that reward can evoke VPL. However, the mechanisms of how reward evoke VPL remain unknown. One possible hypothesis is that VPL is obtained through reward related reinforcement processing. If this hypothesis is true, learning can only occur when reward follows the stimulus presentation. Another interpretation is that VPL is acquired through an enhancement of alertness in association with reward. If the alertness hypothesis is true, learning should occur when reward precedes the stimulus presentation. In our study, we tested the plausibility of the two hypotheses by manipulating the order of reward and stimulus presentation. In Experiment 1, we separated participants into two groups. During training, the ‘Before’ group received water reward 400ms prior to the onset of trained orientation stimulus while the ‘After’ group received water reward 400ms subsequent to the onset of trained orientation stimulus. Both groups were trained using the Continuous Flash Suppression paradigm to render the stimulus imperceptible to the participants by the presentation of dynamic noise in the untrained eye. We found training only in the ‘After’ group indicating that reward may evoke learning through reinforcement-like processing. In Experiment 2, we excluded the possibility that alertness may not be sufficient to elicit learning when presented before stimulus. We presented beep sound prior to the onset of stimulus to increase alertness. Our finding demonstrated that alertness is sufficient enough to evoke learning. In conclusion, our study provided evidence that reward can evoke VPL through reinforcement process.

## Introduction

The role of reward in human behavior has been investigated comprehensively since Pavlov’s classical conditioning experiments(Rescorla & Wagner, 1972; Wolfram Schultz, 2006; W. Schultz, Dayan, & Montague, 1997). How reward promotes learning has been a central question. One of the most dominant models indicate that conditioning occurs when a stimulus is predictive of reward (Rescorla & Wagner, 1972). Schultz and his colleagues have found that dopamine neurons in the substantia nigra were activated when rewards occurred at unpredicted times and were depressed when rewards were omitted at the predicted times (Wolfram Schultz, 2002, 2007). They developed the theory in which conditioning is driven by the prediction error between the reward expected and received.

Several studies have shown that reward also plays a critical role in promoting visual perceptual learning (VPL) (Law & Gold, 2008, 2009; Xue, Zhou, & Li, 2015), which is defined as a long-term performance improvement as a result of visual experiences (Dosher & Lu, 2017; Gilbert & Li; Levi, 2012; Li, 2016; Li, Piëch, & Gilbert, 2004; Sagi, 2011; Seitz & Dinse, 2007; Seitz, Kim, & Watanabe, 2009; Watanabe & Sasaki, 2015)

Seitz and his colleagues (Seitz et al., 2009) presented a sequence of two orientations in random order which were made invisible by the continuous flash suppression paradigm (Tsuchiya & Koch, 2004) One orientation was paired with reward. The other orientation was not paired with reward. All of the orientation stimuli were consistently presented in one eye for each subject. The results showed that performance was enhanced only with the orientation paired with reward and no transfer was found to the untrained eye. These results suggest that reward plays a significant role in VPL, which is involved in early stages of visual processing.

One possible interpretation of these results is that VPL is enhanced by reinforcement processing. It has been found that learning of visual features is contingent on reward probabilities, which is another necessity for reinforcement to occur (Kim, Seitz, & Watanabe, 2015). Arsenault et al (Arsenault, Nelissen, Jarraya, & Vanduffel, 2013) has also reported a reward modulated decrease in BOLD signal in primate early visual cortex. Law & Gold found that response changes in the lateral intraparietal cortex (LIP) of monkeys fits well with VPL driven by prediction errors (Law & Gold, 2009). These and other studies are in accord with the reinforcement-driven VPL hypothesis.

However, another possible interpretation is that reward increases alertness, which enhances internal signals of a presented stimulus and leads to VPL. It has been suggested that alertness can gate the processing of high priority information (Posner & Petersen). Reward increases the release of norepinephrine which in turn increases alertness, and therefore facilitates visual processing (Aston-Jones & Cohen, 2005; Dinse, Ragert, Pleger, Schwenkreis, & Tegenthoff, 2003). Specifically, stimulus paired with reward can evoke an increase in local field potentials driven by enhanced attention(Frankó, Seitz, & Vogels, 2010). Increased alertness measured as dilated pupil diameters have facilitated visual contrast perception(Kim, Lokey, & Ling, 2017). It is noticeable that the above-mentioned Seitz study(Seitz et al., 2009), the last 100msec of an orientation presentation (500msec) was overlapped with reward delivery. This further increases the possibility that reward enhanced an alertness level which directly enhanced stimulus signals.

Although there have been proposals for both the reinforcement and the alertness hypotheses in association with the mechanisms of reward, there is a lack of evidence that indicate which of the two hypotheses is true. In this study, we examined the underlying mechanism of how reward evokes learning in lower-level visual feature processing with the task-irrelevant learning paradigm. If the reinforcement hypothesis stands, for a stimulus to be learned, the stimulus needs to be presented before reward so that the stimulus is predictive of reward. On the other hand, if the alertness hypothesis stands, learning should occur when reward precedes the stimulus. To address the question as to which hypothesis is true, we conducted an experiment in which, a stimulus was presented before or after reward. We found that VPL occurred when a stimulus was presented before reward, but VPL was not observed when it preceded reward. We further found that VPL was not observed with the eye which is different from the eye in which stimuli were presented during training, even when a stimulus was presented prior to reward. These results rule out the alertness hypothesis and suggest that VPL occurs through interactions between early visual areas and reinforcement processing that originates outside the visual system.

## Results

### Experiment 1

We conducted the first experiment using a typical perceptual learning procedure which consists of 12 training sessions and the pre- and post- test sessions before and after the training, see Figure 1 and Method. During the test sessions, orientation discrimination tasks were performed to measure participants’ sensitivity. During the training sessions, participants were randomly separated into two groups which differed in the order of the presentations of reward and stimuli. We presented a sequence of two oriented gratings only in the trained eye in both groups. The Reward Before Stimulus (‘Before’) Group received water reward 400 ms before the onset of the trained orientation presentation. The Reward After Stimulus (‘After’) Group received water reward 400ms after the onset of the trained orientation presentation. No reward was given accompanying the untrained orientation presentation. Moreover, we used the Continuous Flash Suppression paradigm (Tsuchiya & Koch, 2004) in which a continuous sequence of high-contrast, contour-rich random patterns were presented to the untrained eye to render the less salient stimulus in the trained eye imperceptible. The Continuous Flash Suppression paradigm has been demonstrated as a robust paradigm that can cause invisibility of the stimulus in the suppressed eye. The suppression effect can be reliably achieved from the onset of the stimulus to avoid potential breakthroughs. Moreover, the effects of CFS is less susceptible to the eye movement of the participants (Yang, Brascamp, Kang, & Blake, 2014). After the posttest, we conducted an awareness test and asked participants to indicate if they see an orientated stimulus pattern and the orientation of it by pressing the corresponding button. The stimulus was presented using the same paradigm as in training.

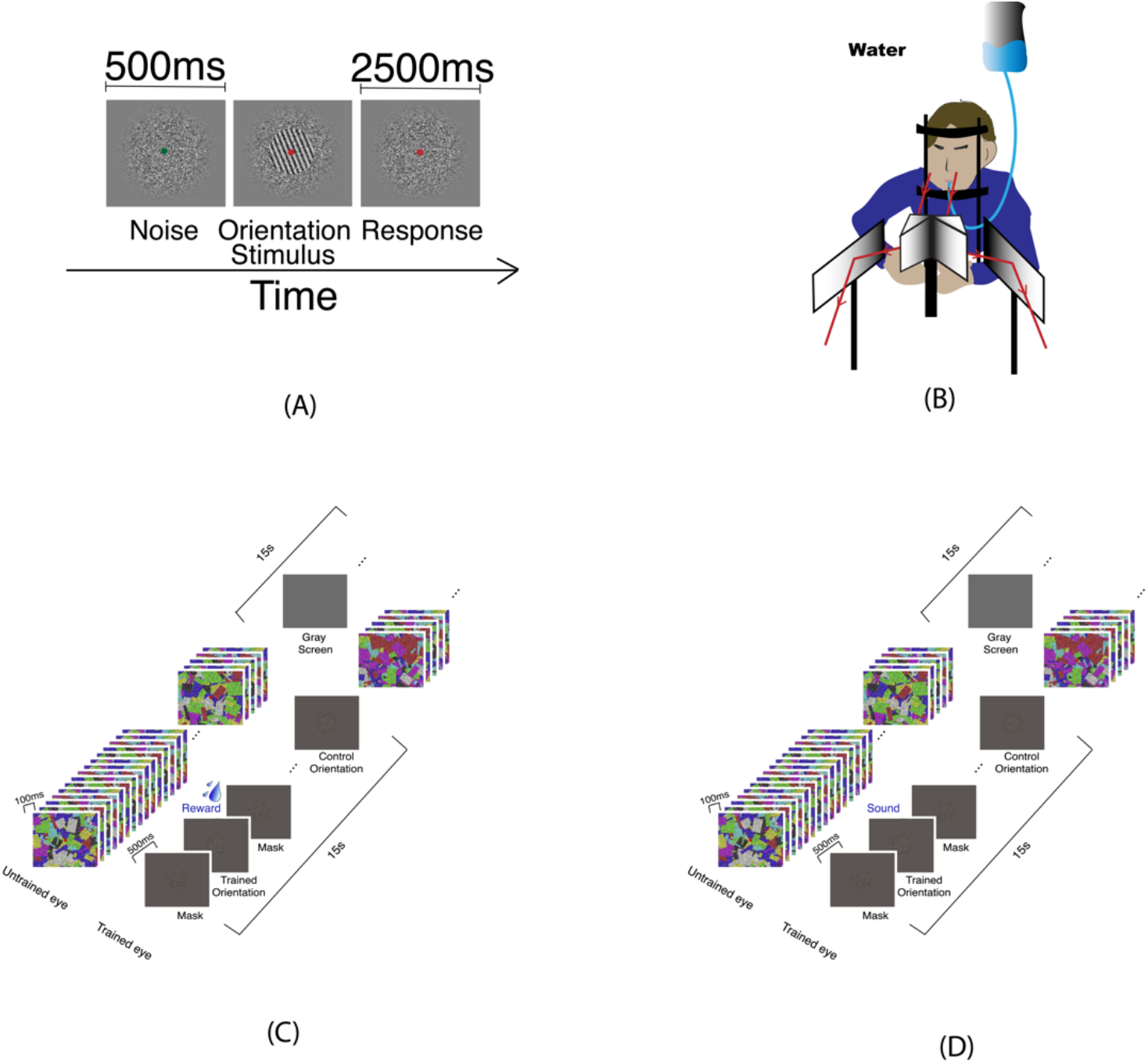

Performance for the orientation discrimination task was characterized by the percentage of correct responses under different S/Ns during pre- and post-test. The percent correct for two orientations in the two eyes were shown in separate panels of Figure 2 for two groups ‘Before’ and ‘After’ respectively. We calculated the improvement in performance as (Pretest-Posttest)/Pretest. A three-way mixed model ANOVA was performed for improvement with ‘Orientation’, ‘S/N ratio’ as the within-group effect and ‘Group’ as the between-group effect. We found a significant significant interaction of ‘Group’ x ‘Orientations’, *F* (3,48) = 3.46, *p*<0.05. To rule out the fact that, the improvement was a result of difference in pre-test performance, we performed a separate ANOVA for the pretest performance with ‘Group’ as the between-group effect and ‘Orientation’, ‘S/N’ ratio as the within group effect. We didn’t find a significant effect of the ‘Group’, *F*(1, 16) = 0.458, *p*>.05 or ‘Orientation’ *F*(3, 48) = 2.64, *p*>.05, or the interaction between them, *F*(3, 48) = 0.513, *p*>.05. We only found a significant effect of ‘S/N ratio’ in pretest performance, *F*(6, 96) = 0.513, *p*<.001. we performed separate ANOVA tests for the ‘After’ group and the ‘Before’ group with ‘Orientation X S/N ratio’ as the main effect, we didn’t find a significant difference in the pretest performance for different Orientations, *F*(3, 24) = 2.353, *p*>.05 for the ‘After’ group and *F*(3, 24) = 0.932, *p*>.05 for the ‘Before’ group.

**Figure 2.**
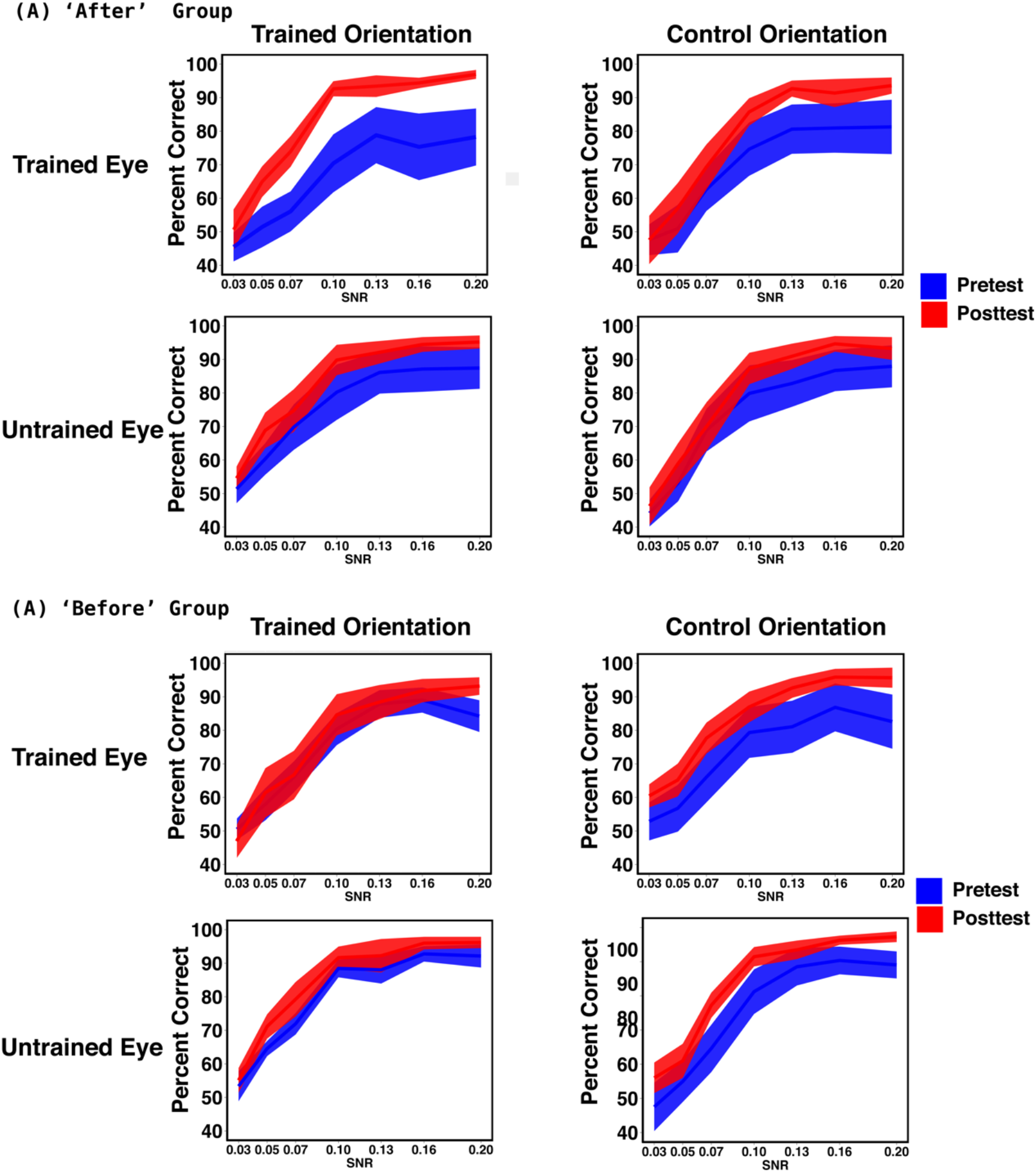
Result of Experiment 1. (A) Result for ‘After’ Group. Precent corret for Pretest and Post test measured for trained orienation and control orientaition in the trained eye and untrained eye, Improvement was only found in the trained orientation in the trained eye. (B) Result for ‘Befor’ group Percent corret for Pretest and Posttest measured for trained orientation and control orientation in the trained and untrained eye. No significant improvement was found.

We performed repeated measures ANOVA in the ‘After’ group and the ‘Before’ group separately with ‘S/N ratio’ and ‘Orientation’ as the within-subject factors. In the ‘After’ group, we found a marginal significant effect of ‘Orientation’, *F*(3, 24) = 2.404, *p*<.01. There was no significant effect in the ‘Before’ group.

In the ‘After’ group, post-hoc t tests showed that there was an improvement significantly different from 0 in the trained eye and in the trained orientation specifically. At the 0.07 S/N level, there was a significant improvement in percent correct after training with *t* (8) = 2.485, *p*<.05. There was no significant improvement in the control orientations and in the untrained eyes, all *p*s >.05. In the ‘Before’ group, we found a slight 10% improvement in the trained orientation in the trained eye and the control orientation in the untrained eye only at the 0.2 S/N level. No significant improvement was observed in the other conditions. As the stimuli at 0.2 S/N level should be clear and conspicuous to the participants, the improvement in the 0.2 S/N level could be a result of improvement in performing the task itself.

In both groups during the awareness tests, participants rarely pressed button and reported that they didn’t see any oriented pattern. In the participants who responded, the accuracy was 0.95% ±2% for the ‘After’ group and 2.78% ± 8.33% for the ‘Before’ group.

To rule out the possibility that the improvement was a result of bias in response, we analyzed the d’ for both groups in the trained eye and the untrained eye. The d’ for both eyes were shown in Figure 3 for two groups ‘Before’ and ‘After’ respectively. Three-Way mixed model ANOVA was performed with ‘Group’ as the between-group effect and ‘Eye’, ‘S/N ratio’ as the within group effects. We found significant ‘Group’ main effect, *F*(1,112) = 10.626, *p*<.01. We also found significant ‘Eye’ X ‘Group’ interaction, *F*(1,112) = 5.179, *p*<.05. We also found a significant interaction of ‘Eye’ X ‘S/N ratio’, *F*(6,48) = 3.345, *p*<.01. Post-hoc analysis indicated that there is a significant improvement in trained eye at 0.1 S/N ratio level, *t*(7) = 3.074, *p*<.05 and the 0.16 S/N ratio level, *t*(7) = 2.39, *p*<.05.

**Figure 3.**
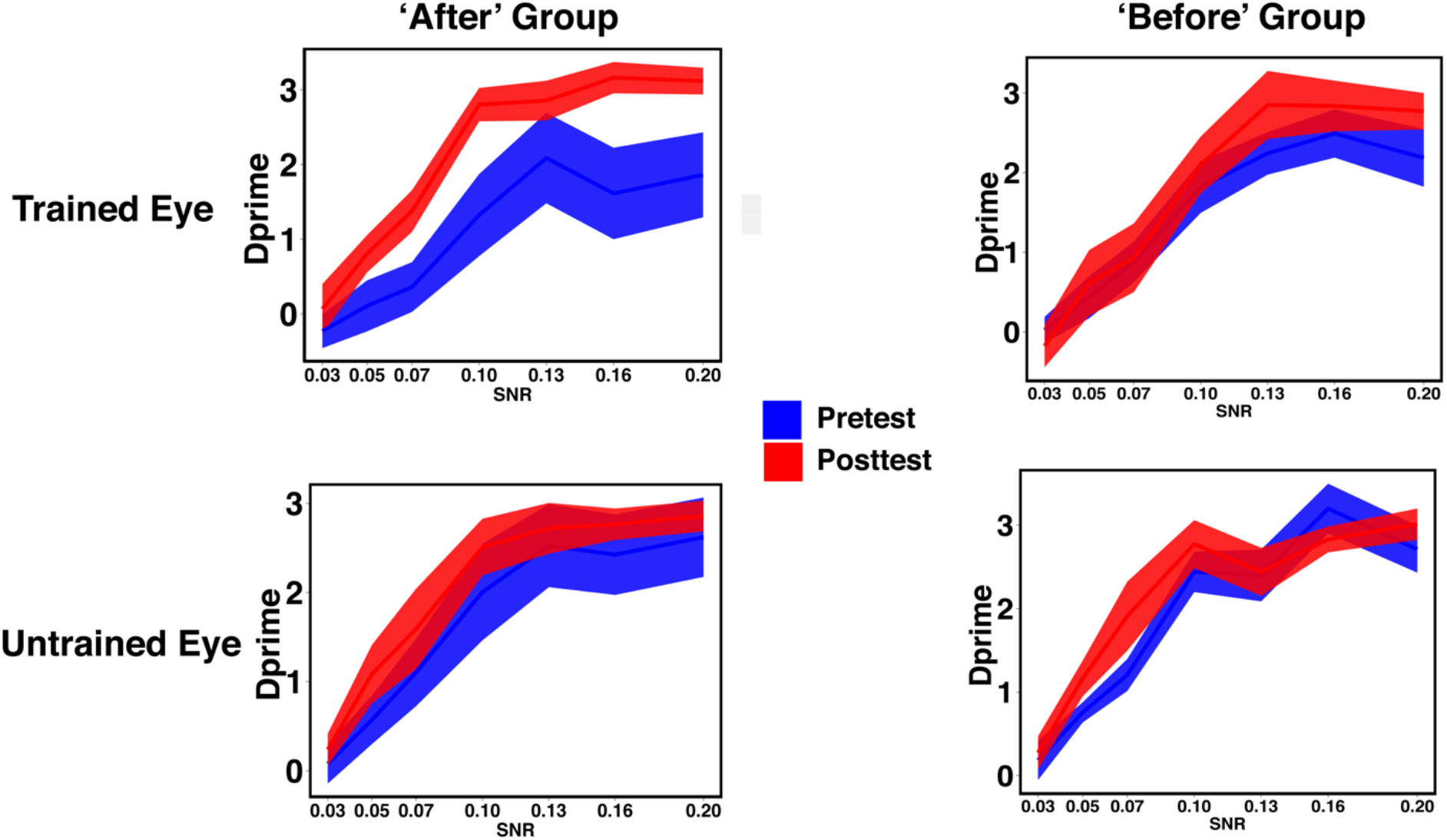
d’for Experiment 1 in two groups obtained with the trained eye and the untrained eye. There is a significant main effect of ‘Group’, *F*(1, 112) = 10.626, p<.01. and a significiant ‘Group’ x ‘Eye’ interaction, *F*(1, 112) = 5.179, *p*<.05. Post-hoc analysis showed significant improvement in d’in the traned eye of the ‘After’ group.

### Experiment 2

There were at least two possible reasons why VPL of a visual stimulus did not occur when reward preceded the stimulus in Experiment 1. The first is that VPL is driven by reinforcement processing but not alertness enhancement. The second is that like reward, somehow alertness needs to be enhanced after a stimulus for VPL. To test which possibility is more likely, we conducted Experiment 2 in which we presented a beep, instead of reward, to enhance alertness before stimulus presentation with the otherwise identical methods to those for The Reward Before Stimulus (‘Before’) Group in Experiment 1.

As in Experiment 1, performance during pre- and post-test was characterized as percentage of correct responses for orientation discrimination task under different S/N ratios. Percent correct for two orientations and two eyes were shown in separate panels in Figure 3. A two-way repeated measures ANOVA was performed with ‘Orientation’, ‘S/N ratio’ as the within-group effects. We found a significant main effect of ‘S/N ratio’, *F*(6,42)=3.18, *p*<0.05. The main effect of ‘Orientation’, *F*(3, 21)=0.601, *p*> 0.05 and the interaction was not significant, *F*(18,126)=0.821, *p*>0.05. Post-hoc analysis showed a significant improvement in percent correct for the orientation that was paired with beep in the trained eye at the 0.1 S/N level, *t*(7) = 3.002, *p*<.05. There is also a significant improvement in percent correct for the orientation that was paired with beep in the untrained eye at the 0.1 and 0.13 S/N level, *t*(7) =2.88, *p*<.05 and *t*(7) = 2.38, *p*<.05 respectively. A significant improvement of the control orientation in the trained eye was also observed at the 0.05 S/N level, *t*(7) = 2.54, *p*<.05.

As in Experiment 1, d’s for the trained eye and the untrained eye were also analyzed and shown in Figure 5. Two-way repeated measures ANOVA with ‘Eye’ and ‘S/N’ ratio as the within-subject effects was performed. We found a significant main effect of the ‘S/N’ ratio, *F*(6,42) = 3.012, *p*<.05. Post-hoc analysis showed a significant improvement of the d’ in the trained eye at 0.1 S/N ratio level, *t*(7) = 2.54, *p*<.05 and significant improvement of the d’ in the untrained eye at 0.1 S/N ratio level *t*(7) = 2.73, *p*<.05 and 0.13 S/N ratio level, *t*(7) = 4.03, *p*<.01.

**Figure 4.**
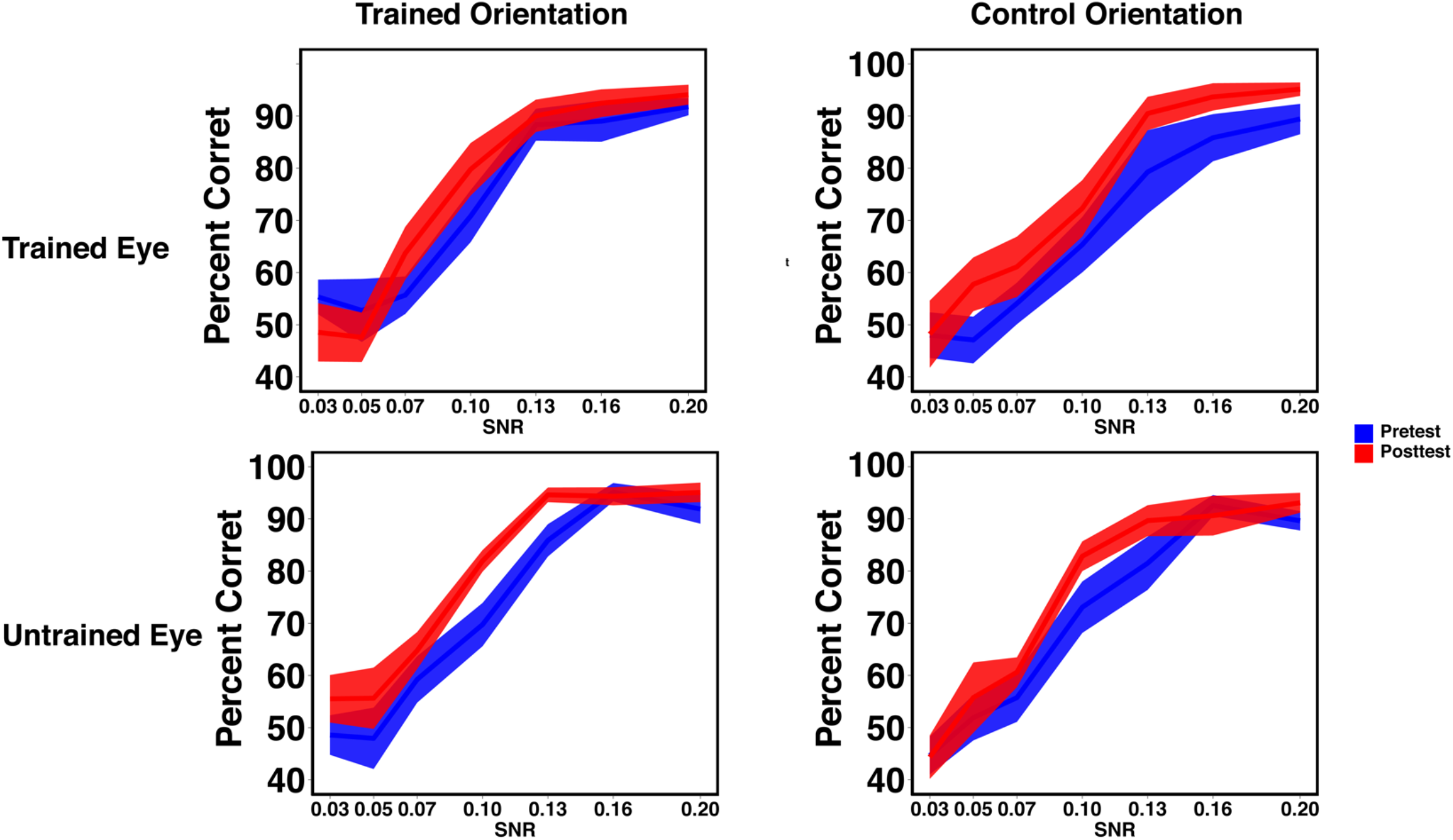
Result of Experiment 2. Percent correct for Pretest and Posttest measured for trained orienation and control orientation in the trained eye and untrained eye.

**Figure 5.**
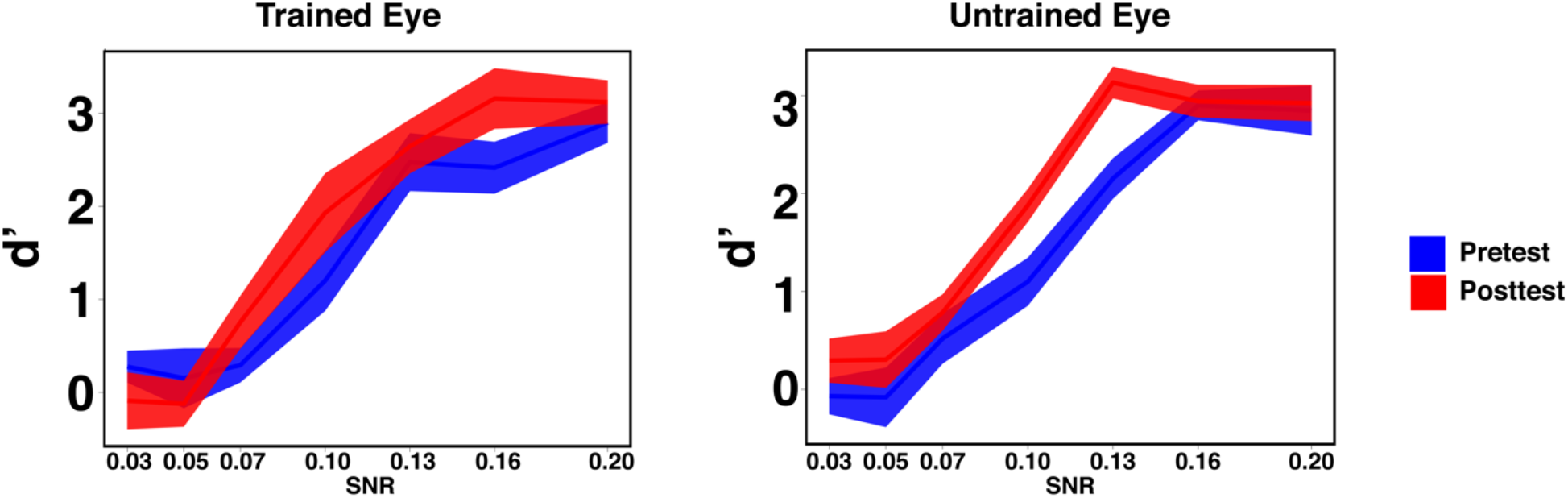
d’of Experiment 2. There was a significant main effect of ‘S/N ratio’, *F*(6,42) = 3.012, *p*<.05. Post-hoc analysis showed that there was significant improvement in d’in both the trained and un-trained eye.

Awareness tests were also conducted as in Experiment 1. The accuracy for participants who responded during the awareness tests were 0.067 ± 0.14.

The results indicate that unlike reward alertness enhancement leads to VPL of a stimulus even if alertness is enhanced before the stimulus presentation. This rules out the possibility that alertness enhancement leads to VPL of a stimulus only when it occurs after the stimulus presentation. Therefore, we conclude that VPL is driven by reinforcement processing.

## Discussion

Our study was aimed to resolve the previous controversy surrounding the role of reward in visual perceptual learning. The reinforcement hypothesis indicates learning should occur only when reward is given subsequently to a stimulus, while according to the alertness hypothesis learning is promoted even when reward precedes the stimulus. In Experiment 1, we conducted two conditions; in the reinforcement condition one of the two orientations that were presented in sequence was presented prior to reward and in the alertness condition it preceded reward. The results demonstrated that VPL of the orientation occurred only when the stimulus preceded reward. This is in accordance with the reinforcement learning hypothesis. Furthermore, VPL was restricted to the trained eye, which suggests that early visual areas are associated with reinforcement-driven VPL. We also conducted the Experiment 2 in which one of the two orientations was presented with a beep sound prior to it. We found improvement in the orientation that was paired with sound which rules out the possibility that alertness before stimulus is not sufficient for performance improvement.

Previous evidence demonstrated that reward evokes both reinforcement signals (Wolfram Schultz, 2006) and alertness (Aston-Jones & Cohen, 2005). Nevertheless, it seems that at least in the current experimental setting VPL is driven by reinforcement processing but not by alertness. In our study, we attempted to increase the possibility of VPL by introducing a 100ms overlap between the reward and stimulus presentation. However, VPL did not occur despite the overlapping interval in the reward before stimulus group. Therefore, the contiguity requirement-reward following the stimulus- seems to be a prerequisite for reward elicited learning to occur.

When alertness is directly manipulated through sound, we not only found improvement of the trained orientation in the trained eye, we also observed improvement transferred to the untrained eye. Alertness might increase along with the changes in attention. The transfer of learning may occur as a result of involvement of higher-order processing in the brain as a result of increase in alertness.

Although decision areas are typically associated with reinforcement learning, our results that VPL did not transfer to the untrained eye suggests the involvement of early visual areas in reinforcement driven VPL. This is consistent with studies in which reward/ reinforcement processing changes activity in the primary visual cortex. It has been found that reward induces reduced signal in the early visual cortex when reward is predictive of a visual cue (Arsenault et al., 2013). Neurons in the rodents’ primary visual cortex learned to be activated to the stimulus that predict reward (Shuler & Bear, 2006). In addition to the findings with animals, the improvement in d’ in our results indicates that the it is not induced by a change in response bias but rather a sensitivity enhancement to the trained features. All of these results suggest that reinforcement processing reaches early visual areas which involved VPL.

A number of studies have indicated that reward plays a crucial role in VPL. However, it remained unclear whether reinforcement processing or alertness caused by reward is a major factor on VPL. We conducted experiments in which the temporal order of the presentations of a visual stimulus and reward was varied. We found that VPL of the stimulus occurs when the visual stimulus was presented before reward but not when it followed reward. In addition, such reward-driven VPL did not transfer to the untrained eye. These results suggest that it is reinforcement processing and not alertness which interacts with visual stimulus signals in early visual areas and leads to VPL.

## Method

### Participants

A total of 26 participants were recruited in this study. 18 participants (13 females) were recruited for Experiment 1 and 8 (6 females) participants were recruited for Experiment 2. Participants were adults (aged between 18 and 60) with normal or corrected-to-normal vision. This study was approved by the Institutional Review Board (IRB) of Brown University. Written consent forms were provided to participants in accordance with the IRB. Participants had never participated in visual training experiments prior to this study.

### Stimuli

Noise masked sinusoidal gratings were presented during test and training sessions. The gratings were orientated 112.5° or 22.5°, with 4° diameter, 10% contrast and a spatial frequency of 2 cycle/degree. The gratings were presented at the center of an 8° annulus surrounded by gray background. The 4° to 8° field of the annulus was presented with Gaussian masked random noise. The stimuli were presented at the center 0° to 4° masked by noise. The noise was generated from a sinusoidal luminance distribution, which ensured that the statistical distributions of the luminance for the noise and the gratings were identical to each other. Consequently, there were no texture cues associated with different levels of noises. During test sessions, 7 different signal-to-noise levels (SN: 0.03, 0.05, 0.07, 0.1, 0.13, 0.16 and 0.2) were varied from trial to trial. During training sessions, a constant SN level of 0.2 were used.

## Experiment 1

### Pretest and Posttest

Sensitivity tests were performed during Pretest and Posttest sessions. These test sessions were scheduled at least one day apart from the training sessions. During the test sessions, stimuli was presented to one eye and gray screen was presented with the other eye. Participants were instructed to align the two screens before starting each testing block. At the beginning of each trial, random noise was presented with a green fixation point for 500ms. The noise was followed by the grating stimuli and a red fixation point for 500ms. The red fixation point will be presented for another 2500ms after the stimuli disappeared. The red fixation point indicated the presence of stimuli as well as the signal for participants to make responses. Participants were instructed to judge whether 112.5° or 22.5° grating was presented by pressing the corresponding buttons. 7 SN levels was presented for each of the two orientations in two eyes separately. The 28 conditions were pseudorandomly interleaved with 32 trials for each condition. Participants were tested for a total of 896 trials in total during the pretest and posttest sessions.

### Training

There were twelve training sessions. Each training session took place on different days. Participants were asked to abstain from eating and drinking for five hours before each training session. Water was provided for reward to the participants during the training period. Participants were separated into two groups --Reward Before Stimulus Group and Reward After Stimulus Group – differed in the timing between the delivery of reward and the presentation of grating stimuli. Continuous Flash Suppression (CFS) paradigm (Tsuchiya & Koch, 2005) was adopted during the training period to render the stimulus invisible to the participants. This procedure eliminated subject’s conscious bias of associating grating stimuli with the presence or absence of reward. To ensure contiguous suppression from flash patterns, we adopted an alternating mini-block of 15s CFS stimuli to each eye. During the first mini-block, the untrained eye was presented with high contrast flashing noise. The CFS stimuli consisted of a sequence of full-screen textured pattern images presenting at a rate of 10 Hz. The texture pattern consisted of 300 randomly placed, physically overlapping rectangles or ellipses of various sizes with dimension from 0.5° to 5°. The shapes were of different orientations as well as saturated colors (0 or 100 cd/m^2^). In addition, 50% of the screen was covered with spatially sparse, colored noise. The trained eye was presented with 2Hz noise. Intermittently, a grating stimulus (112.5° or 22.5°) of 0.2 SN was presented to the trained eye for 500 ms. One of the orientations was chosen randomly as the trained orientation. The time interval between the grating stimulus was at least 3000 ms. For the Reward Before Stimulus Group, water was delivered 400ms prior to the presentation of the trained grating stimulus. For the Reward After Stimulus Group, water was presented 400ms following the presentation of trained grating stimulus. No reward was paired with the untrained grating stimulus. In both cases, reward was presented for 500ms with 100 ms overlap with the stimulus presentation. The overlap ensures processing of reward and maximizes the possibility that water reward could be associated with orientation processing. After the first 15s mini-block, the trained eye was presented with the CFS stimuli while the untrained eye was presented with a gray screen. No reward was presented during this mini-block. The 15s mini-blocks alternated for 5 minutes, and participants were asked to take a 3-minute break after the 5-minute block. These sequences were repeated for 6 times per session, yielding a total of 120 trials for the trained and untrained orientations respectively. Participants were requested to adjust the haploscope to align the two screens after each break.

### Awareness Test

Awareness test was conducted immediately after the Posttest to ensure the CFS paradigm has successfully rendered the stimuli invisible. Participants were presented with the same stimuli as in training sessions for one block. During the awareness test, participants were asked to decide whether there was a grating presented and the orientation of the grating by pressing corresponding buttons. Participants were also interviewed whether they were able to detect the grating during the awareness test session as well as during the previous training sessions. Participants were also inquired about whether they were aware of any patterns in the timing of water delivery.

## Experiment 2

### Pretest and Posttest

Participants performed the same sensitivity tests during pre- and post-test as in Experiment 1.

### Training

Participants were trained for 12 sessions. The procedure was the same as the Reward Before Stimulus Group in Experiment 1 except that a beep sound was presented 400ms before the stimulus presentation instead of reward. Additionally, participants were not instructed to abstain from eating and drinking before each training session.

### Awareness Test

The procedure for Awareness tests was the same as in Experiment 1.

### Apparatus

Stimuli were presented on two 19□□ CRT screens (1024 × 768 pixels, 120 Hz). The luminance of the screen was gamma corrected. The stimuli were presented through MATLAB (The MathWorks, Natick, MA) and Psychtoolbox-3 (Brainard, 1997; Pelli, 1997; Kleiner et al., 2007) using a Mac OSX computer. Participants viewed the stimuli through a haploscope. The viewing distance was 82cm. The water delivery equipment was ValveLink→8.2 system manufactured by Automate Scientific, Inc.

## Acknowledgements

This study was supported by NIH R01EY019466 and NSF BCS 1539717 to Yuka Sasaki.

## Author Contributions

Z.W., Y.S. and T.W. designed the study. Z.W. and D.K. wrote the experimental codes. Z.W. conducted the experiments and analyzed the data. Z.W., Y.S. and T.W. wrote the manuscript.

